# Incorporating Large Language Model-Derived Information into Hypothesis Testing for Genomics

**DOI:** 10.1101/2025.04.30.651464

**Authors:** Jordan G. Bryan, Hongqian Niu, Didong Li

## Abstract

We propose strategies for incorporating the information in large language models (LLMs) into statistical hypothesis tests in genomics studies. Using gene embeddings derived from text inputs to OpenAI’s GPT-3.5 model, we show that biological signals in a variety of genomics datasets reside near the principal subspace spanned by the embeddings. We then use a frequentist and Bayesian (FAB) framework to propose several hypothesis tests that are either optimal or approximately optimal with respect to prior information based on the gene embedding subspace. In four real-world genomics examples, the FAB tests guided by the LLM-derived information achieve more power than classical counterparts.

## 1 Introduction

In a recent review article, Simon et al. (2024) identified exploratory embedding analysis, prediction, and transfer learning as key uses of LLM-derived information for discovery in the biological sciences. In this article, we demonstrate that statistical inference may be added to the list of tasks for which LLM-derived information may improve discovery in genomics studies.

In the past decade, foundation models based on deep convolutional neural networks such as ResNet (He et al., 2016) or large transformer networks such as Google’s BERT (Kenton and Toutanova, 2019) and OpenAI’s GPT family (Radford, 2018) have revolutionized the analysis of data across fields such as computer vision and natural language processing. In addition to their commercial applications, these models have been shown to be valuable aids in scientific discovery in areas ranging from language understanding and translation (Vaswani et al., 2023) to multi-modal biomedical applications such as medical imaging (Li et al., 2024), drug prediction (Zhang et al., 2024), and analysis of genomic data (Cui et al., 2024; Theodoris et al., 2023; Chen and Zou, 2023).

Foundation models for genomics applications can be roughly divided into two categories: those that are trained on enormous corpora of experimental data and those that are trained on internet-scale databases of natural language. Models in the second category may be trained on databases filtered to include only biology-related texts (Lee et al., 2020; Gu et al., 2022), or not (Nori et al., 2023). Models that fall into the first category include the scGPT model (Cui et al., 2024), which was trained on transcriptomes from 33 million human cells from 441 different studies, curated from the CellxGene Collection (Program et al., 2024) and the GeneFormer model (Theodoris et al., 2023), which was trained on a dataset comprised of 29.9 million human single-cell transcriptomes denoted Genecorpus-30M, collated from over 561 publicly available datasets.

While it may seem natural to construct a foundation model for biology from biological data, models that fall in the natural language category have some distinct advantages. For instance, high-quality text data are more abundant than high-quality sequencing data. Perhaps more importantly, there may be relationships between biological entities that are well-documented in the scientific literature, but which are not present in large-scale genomics datasets because they arise only in specific contexts that require bespoke experimental designs. For this reason, some recent works have used the embedding outputs of large-language models (LLMs) such as ChatGPT (Radford, 2018) to encode the biological information contained in text-based gene descriptions, such as those in the NCBI database (Schoch et al., 2020). Notably, Chen and Zou (2023) show that these text-based gene descriptors can be input to GPT-3.5 to obtain gene embeddings that act as covariates for standard prediction algorithms. For various gene-level tasks such as gene functionality class prediction, gene property prediction, and gene-gene interaction prediction, using these gene embeddings as covariates for a random forest algorithm was shown to have favorable predictive performance, even compared to that of specially pre-trained transformer models such as scGPT and GeneFormer, or BiolinkBERT (Yasunaga et al., 2022). Other works in this direction include Hou and Ji (2024), who demonstrate that GPT-4 is capable of generating cell type annotations from differential marker genes, Märtens et al. (2024) who show that incorporating prior knowledge through LLM-informed gene embeddings can achieve state-of-the-art performance on predicting unseen perturbation-induced transcriptomic changes, or Hu et al. (2024) who demonstrate the potential for pre-trained LLMs to identify gene set functions.

Based on this body of research, it is evident that LLMs are capable of producing numerical embeddings of biological entities such as genes, which capture information with relevance to a number of prediction tasks. Less studied to date is the potential for LLM embeddings to guide statistical inference in genomics contexts. In this article, we document some strategies for incorporating LLM-derived information into statistical tests of certain hypotheses that commonly arise in genomics studies. We propose several hypothesis tests, which have exact or approximate optimality properties with respect to the gene-to-gene similarity information encoded by LLM gene embeddings. While the tests are based on different parametric sampling models, all of them are motivated by the supposition that biologically meaningful signals reside on or near a subspace determined by the major axes of variation in LLM gene embeddings. Importantly, each test maintains type I error guarantees regardless of whether that supposition is accurate.

### A case study in basal mRNA expression of blood cancers

We explore what it means for a biological signal to be close to a subspace determined by LLM gene embeddings through an example involving a subset of mRNA expression data downloaded from the Cancer Dependency Map (DepMap, 2024). The subset of gene expression measurements we consider consists of 728 Cancer Gene Census genes (Sondka et al., 2018) measured in 31 acute myeloid leukimia (AML) cancer cell lines and 24 B-cell acute lymphoblastic leukimia (BLL) cancer cell lines. We compute the average expression profiles for the AML and BLL cancer cell lines, and then take the difference between these averages to be a biological signal of interest. Along with the gene expression data, we assemble a matrix of LLM-derived gene embeddings, **E** ∈ ℝ^728*×*1536^ downloaded from the GenePT repository (Chen and Zou, 2023).

Let 𝒱^728^ denote the set of 728 *×* 728 orthogonal matrices, let **U** ∈ 𝒱^728^ be the orthogonal matrix of left singular vectors of **E**, and let **y** ∈ ℝ^728^ denote the signal vector whose entries are the differences in average mRNA expression between the AML and BLL cell lines. For each *k* ∈ *{*1, …, 728*}*, we evaluate the quantity

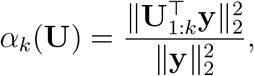

where **U**_1:*k*_ denotes the matrix consisting of the first *k* columns of **U**. This is equal to the norm of the signal vector lying along the first *k* principal axes determined by the gene embeddings, relative to the total signal norm. In Figure 1, we plot *α*_*k*_(**U**) alongside 100 realizations of the random object *α*_*k*_(**V**) for **V** simulated from the uniform distribution on 𝒱^728^. In the right panel of Figure 1, we observe that the first several singular vectors of **E** capture more of the signal norm than would be expected by random projection, since the dashed lines indicate the 2.5% and 97.5% quantiles of the realizations of *α*_*k*_(**V**).

**Figure 1.**
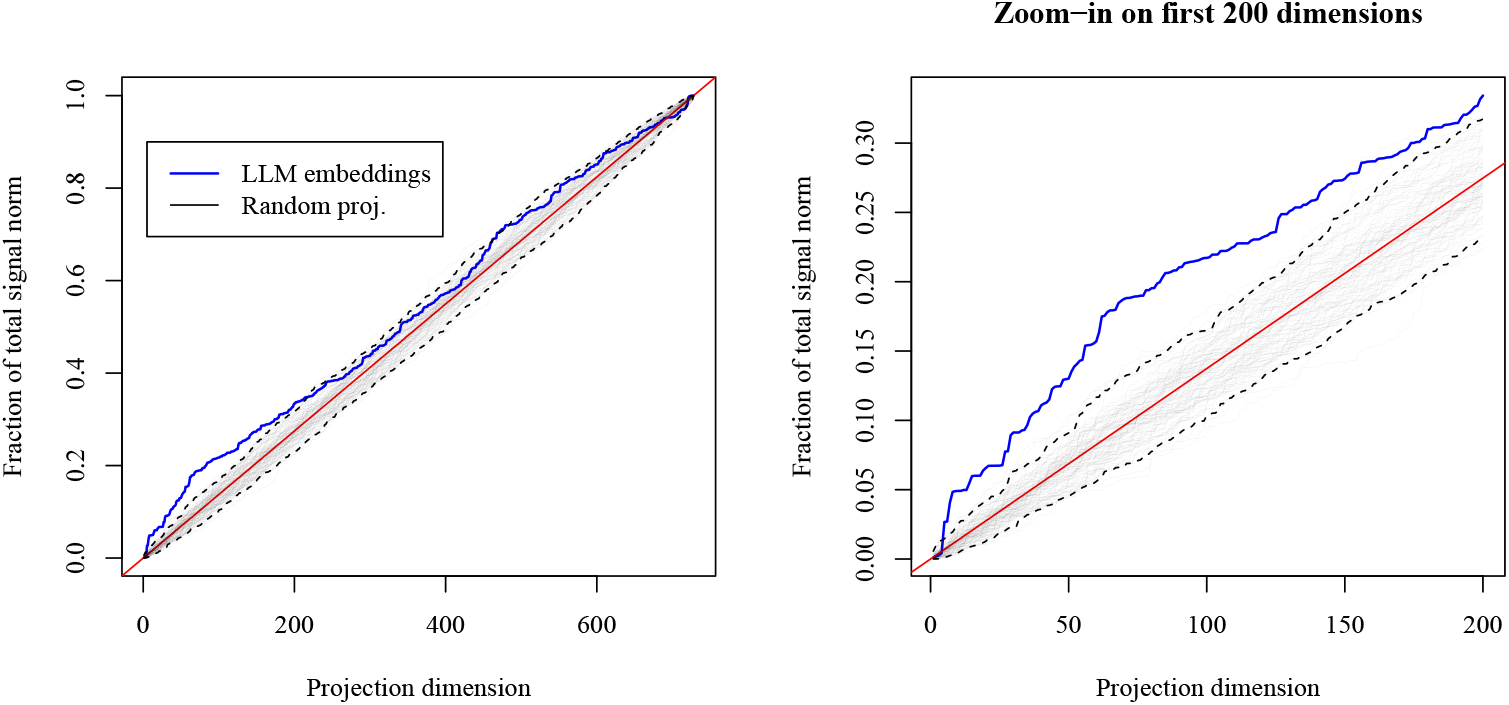
The quantity *α*_*k*_(**U**) versus projection dimension *k*, for *k* ∈ {1, …, 728} . LLM embeddings appear in blue, while random realizations of *α*_*k*_(**V**) appear as thin, transparent lines, with dashed lines as 2.5% and 97.5% quantiles.

The results of this small experiment both confirm long-standing intuition about the nature of biological signals with gene-level measurements and also indicate how LLMs might be used to detect them. Two hallmarks of modern genomic data analysis are the concepts of dimension reduction and gene-set-level analysis. Both of these approaches are motivated by the belief that in a *p*-dimensional measurement space where each coordinate corresponds to a gene, biologically relevant results will not appear uniformly at random within the space, but will rather tend to appear near low-dimensional subspaces that are determined by how genes relate to each other in a given experimental context. The LLM-derived gene embeddings we used in this experiment were not designed to span or approximate any specific subspace pertaining to genes relevant to AML, BLL, or the difference between these two cancer subtypes. Rather, it appears that the gene embeddings encode a general-purpose notion of gene-gene similarity, which—while not perfectly tuned for any specific experiment—may nonetheless provide some value for detecting biological signals in a variety of experimental contexts.

In this article, we translate the general task of detecting biological signals into specific statistical hypothesis testing problems that arise in genomics studies. We then provide a means of incorporating the information in LLM-derived gene embeddings into the evaluation of each hypothesis. The methodology we propose is motivated by using the LLM information to specify a Bayesian prior distribution over the parameter that indexes the null hypothesis for each test. However, the proposed hypothesis tests are assessed using the frequentist paradigm and are designed to maintain type I error control at a desired level. Hence, they are termed “frequentist and Bayesian” (FAB) (Hoff, 2023).

The primary contributions of this article are two-fold: first, we expand the scope of FAB testing in genomics studies by applying existing FAB tests to new contexts and by proposing two new FAB hypothesis tests, one for multivariate means and one for coefficient estimates from multiple logistic regressions. Next, we use these tests to demonstrate how the predictive power of gene embeddings derived from LLMs can be leveraged for statistical inference. Our goal is to offer more powerful alternatives to classical statistical tests in a variety of realistic analysis scenarios in genomics studies and to couple these tests with a source of information that can be easily accessed by practitioners. In Section 2, we provide a high-level description of how LLM-informed FAB hypothesis testing works. We then outline specific scenarios in the analysis of genomics data where such tests may be applied and we perform experiments comparing results obtained from LLM-informed FAB testing to the results one would obtain using classical statistical tests. We find that LLM-informed FAB tests yield as many or more discoveries for a fixed false discovery rate (FDR) than analogous classical hypothesis tests that do not use external information. In Section 3, we conduct simulation studies that illustrate the type I error control of the FAB tests we propose. We provide full descriptions of the FAB tests used for each analysis scenario in the Methods section. We summarize our contribution and discuss directions for future work in Section 4.

## 2 LLM-informed FAB testing in genomics

In this section, we consider four testing scenarios in modern genomics studies and demonstrate that the FAB tests applicable to each scenario yield as many or more discoveries than do classical tests that do not make use of the LLM-derived information.

### 2.1 Frequentist and Bayesian (FAB) testing

The FAB testing methodology was first introduced by Hoff (2020) who focused on information-assisted modifications of *t*-tests. Bryan and Hoff (2022) later applied these *t*-tests to certain genomics contexts using external information derived from large genomics datasets. More recently, McCormack and Hoff (2023) proposed FAB tests for linear hypotheses, and Bersson and Hoff (2024a,b) used FAB principles to reduce interval size in conformal prediction. While all of these articles—and the present article—make comparisons between FAB procedures and analogous classical statistical procedures, it is important to state that FAB hypothesis tests are entirely consistent with much of the classical frequentist testing paradigm. Most importantly, FAB tests are designed to have the same type I error control offered by frequentist hypothesis tests. The key distinction lies in the notion of bias as it applies to statistical hypothesis testing and in the implications this has for type II error. FAB tests are not necessarily unbiased, while classical tests tend to be. By sacrificing unbiasedness, FAB tests may have improved type II error, i.e. more power, relative to classical tests.

The normal mean testing problem is a valuable example for illustrating these concepts. Suppose that *n* experimental replicates are generated independently according to

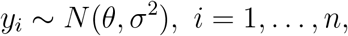

where for simplicity we assume that *σ*^2^ is known. Frequentist statistical theory suggests that an optimal level-*α* unbiased test of the hypothesis *H* : *θ* = 0 versus the alternative *K* : *θ* ≠ 0 is one that rejects *H*_0_ when the statistic 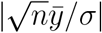 exceeds its 1 − *α* quantile under the null distribution 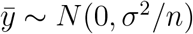. A FAB test of the same hypothesis arises by positing a Bayesian prior distribution over the unknown *θ*

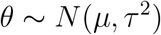

and asking what testing procedure would be optimal at level *α* if that prior information were accurate. In this example, the answer is that a Bayes-optimal level-*α* test results from rejecting *H*_0_ when the statistic 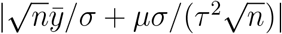 exceeds its 1 − *α* quantile under the null distribution 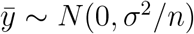. This FAB test statistic includes a shift term, which can be interpreted as the product of two ratios: that of the prior mean to the prior standard deviation and that of the sampling standard deviation to the prior standard deviation. The FAB test based on this statistic may be biased, but its power may be greater than that of the unbiased test if *θ* and *µ* have the same sign.

Figure 2 illustrates the tradeoff in power of the FAB and classical tests for different values of the prior parameter *µ*. In all panels, *n* = 3 and *σ*^2^ = *τ* ^2^ = 1. One way to summarize the display is that the prior mean *µ*—the central point of the densities in the bottom panels—governs a tradeoff between increased power for *θ* values with the same sign as *µ* and decreased power for *θ* values with the opposite sign of *µ*. In fact, it is possible for the power of the FAB test to fall below the level *α* for *θ*’s that have the opposite sign of *µ*; this is what it means for the test to be biased. On the other hand, for *θ*’s that have the same sign as *µ*, the power of the FAB test may be as large as if one knew the sign of *θ* in advance. Notably, none of the tests exceeds the level *α* = 0.05 at the null value *θ* = 0.

**Figure 2.**
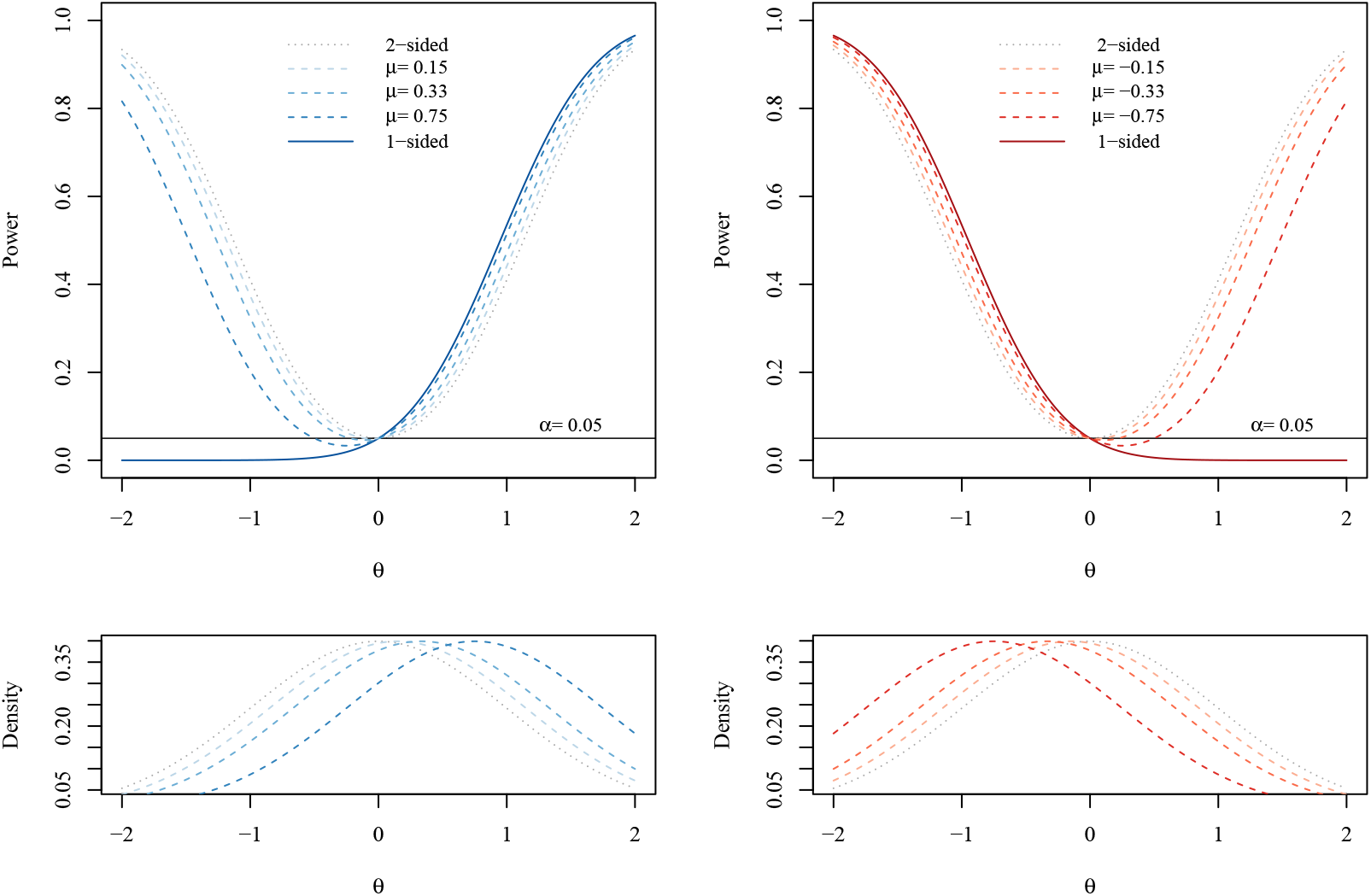
Illustration of the power of six FAB tests compared to one-sided (bold line) and two-sided (gray dotted line) *t*-tests. The top left and right panels display the power curves for positive and negative values of *µ*, respectively. The bottom left and right panels display the prior densities for the different values of *µ*.

Especially in the case of multiple hypothesis tests, a common circumstance in genomics studies, the increase in power afforded by FAB tests informed with accurate prior information can translate into significantly more discoveries than would be made using classical procedures at a fixed false discovery rate (FDR). The genomics data analyses we consider in this article use FAB tests in the context of multiple hypotheses, and they employ LLM-derived gene embeddings as a source of prior information. While the tests we use are designed for more complicated statistical hypotheses than the normal mean problem considered above, the conclusion about how the prior information functions is similar: the power of the FAB tests depends on how informative the LLM-derived information is about the space of realistic biological signals. Among the four real-world genomics applications that follow, we observe both positive and negative results in this regard. However, in the one negative example, the FAB test we employ adapts to the fact that the relevant biological signal does not seem to be captured by the embedding subspace. Therefore, the number of discoveries made for a fixed FDR of 0.1 is still greater, if only modestly greater, than that made by the classical test. In each example below, the LLM information takes the form of a *p ×* 100 matrix of gene embeddings **E**, where *p* is equal to the number of genes under consideration. The embeddings we use are obtained by taking the best rank-100 approximation of the centered and scaled *p ×* 1536 embeddings matrix downloaded from the GenePT repository.

### 2.2 Determination of CRISPRi bioactivity using PerturbSeq

Advances in sequencing technologies have enabled post-perturbation readouts of transcriptional activity to be achieved at genome-scale. In a landmark study, Replogle et al. (2022) used the Perturb-seq screen (Adamson et al., 2016) to record gene expression profiles for the K562 chronic myeloid leukimia (CML) cancer cell line in response to over 10000 perturbations achieved through the CRISPR interference (CRISPRi) technology. Each perturbation in this study corresponds to transcriptional suppression of a certain gene, which may or may not induce a change in the post-perturbation transcriptional state of the cell line. For downstream analyses, it is of interest to determine which perturbations show evidence of inducing some transcriptional change in K562.

To address this question, we applied the two-sample AFAB test for multivariate means described in Section 5.2 to the pseudo-bulk RNA expression profiles of Replogle et al. (2022). After filtering for genes that had GenePT embeddings available, each profile consisted of *p* = 7614 measurements of mRNA expression. In total, there were 10673 unique perturbation profiles, as well as 585 profiles arising from non-targeting single-guide RNAs (sgRNAs). For the purpose of the two-sample tests, the non-targeting profiles were used as the control sample (*n*_1_ = 585) and each perturbation profile was taken to be the treatment sample (*n*_2_ = 1), so only the non-targeting profiles were used to compute the sample covariance **S**.

In Figure 3, we plot the 10673 *p*-values after adjustment with the Benjamini-Hochberg procedure (Benjamini and Hochberg, 1995) that we obtained from each of three testing procedures: random projection (RP), naive AFAB (AFAB), and split-sample AFAB (SS AFAB) with *r* = 0.66. The random projections (RP) procedure was performed according to the steps in Lopes et al. (2011) with a projection dimension equal to 100. We observe that projection onto the embedding column space (*T*_AFAB_ with 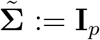) yields a test with more power than the RP test. The split-sample version of *T*_AFAB_ displays a marginal further improvement in power, possibly owing to the fact that the covariance matrix of the non-targeting sgRNA profiles is approximately low-rank (see discussion in Section 3.2).

**Figure 3.**
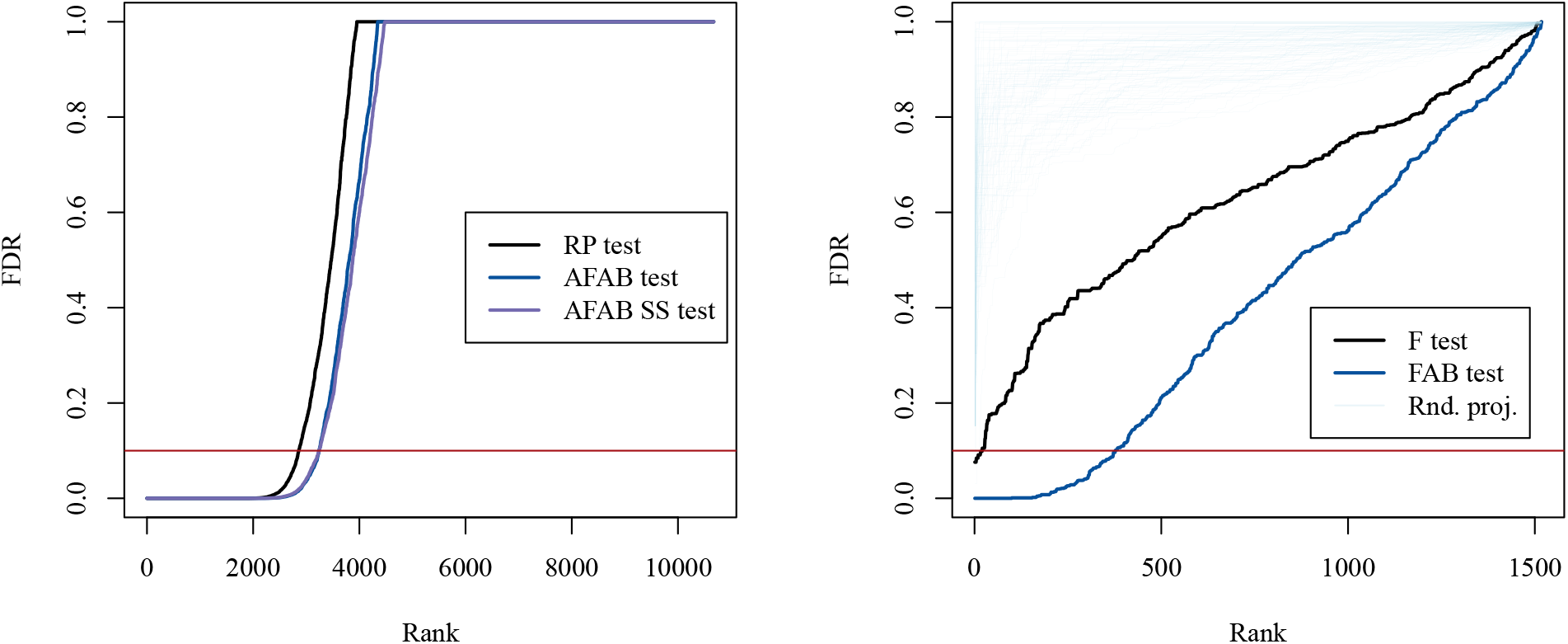
(left) Sorted and adjusted *p*-values from the two sample tests applied to CRISPRi PerturbSeq profiles (Replogle et al., 2022). The naive AFAB test (blue) yields more discoveries than the RP test (black) at an FDR of 0.1. (right) Sorted and adjusted *p*-values from *F* -tests applied to drug discovery data from Corsello et al. (2020). The FAB *F* -test (blue) yields more discoveries at an FDR of 0.1 than the classical *F* -test (black) and random-projection *F* -tests (thin, light blue).

### 2.3 Drug discovery from cell viability profiles

Pharmaceutical companies and research institutions with an interest in discovering new cancer therapies sometimes conduct high-throughput screens to determine the effect that chemical compounds in their vast libraries have on the viability of several cancer types (Broach et al., 1996; Astashkina et al., 2012). The viability effect of a drug on a particular cancer cell line is measured by the relative abundance of a stably expressed transcript in a treated sample of cells with respect to that of a control sample. While an individual viability effect may be of interest in its own right, it is often the case that a compound will be prioritized for further investigation by considering the pattern of its viability effect across multiple cell lines. This viability profile may be compared to transcriptomic data such as baseline mRNA expression in order to determine whether there is evidence of a relationship between the biological characteristics of the cancer cell lines and their response to drug treatment.

We used a cancer viability dataset from Corsello et al. (2020) to investigate whether FAB *F* -tests parameterized by LLM-derived gene embeddings could be useful for drug discovery in the context of a large-scale cancer viability screen. Specifically, we applied the limiting-case FAB *F* -test of Section 5.3 to each of 1518 drug viability profiles, using mRNA expression values from the Dependency Map as the matrix of independent variables **G** ∈ ℝ^*n×p*^. As in Section 1, we considered *p* = 728 genes belonging to the Cancer Gene Census, and after filtering for common cancer cell lines between the expression dataset and the viability dataset, there were *n* = 803 cancer cell lines per viability profile.

We apply both the embedding-projected FAB *F* -test of Section 5.3 and the classical *F* -test to each of the 1518 drug viability profiles and display the results in the right panel of Figure 3. We plot the sorted *p*-values obtained from the FAB and *F* -tests after adjustment with the Benjamini-Hochberg procedure. The classical *F* -test suffers from a lack of statistical power due to the large number of genes under consideration relative to the number of cancer cell lines. Only 20 drugs meet the FDR *<* 0.1 criterion according to the *F* -test, while 193 drugs do so for the FAB test. In light blue, we also plot the adjusted *p*-values corresponding to FAB tests performed using embedding matrices with entries simulated independently from *N* (0, 1). This method of random projection evidently performs worse than the classical *F* -test, demonstrating that the increase in power of the FAB test is achieved not merely through dimension reduction, but through the particular manner of dimension reduction determined by the LLM-derived matrix of gene embeddings.

### 2.4 Analysis of differential dependency in AML

We next consider a differential dependency analysis between 10 Ras-mutant and 8 Ras-wildtype AML cancer cell lines using data from Wang et al. (2017). In this study, CRISPR dependency scores in each AML cell line were recorded for a set of *p* = 132 gene-knockouts comprised of several known Ras interactors. As mutations in the Ras family of genes are common signatures of treatment-resistant cancer (Cox et al., 2014), it is of interest to determine which of the gene-knockouts produce a significant effect on Ras-mutant AML viability relative to that observed in the Ras-wild-type AML cell lines.

To address this question, we applied the FAB methodology of Section 5.4to testing the hypothesis that there is no difference in average viability between Ras-mutant AML and Ras-wild-type AML upon deletion of the *j*th gene for each *j* ∈ *{*1, …, *p}* . For comparison, we also evaluated each of these hypotheses with the classical two-sided *t*-test. As shown in Figure 4, the FAB *t*-tests rejected more null hypotheses than the classical *t*-tests at an FDR of 0.1 after *p*-value adjustment. Interestingly, one of the genes discovered only by the FAB *t*-tests is *KRAS*, a member of the Ras family. Gene-knockout of *KRAS* is known to be preferentially lethal to Ras-mutants (Meyers et al., 2017), so its inclusion in the list of discoveries is to be expected. The genes *GNG5* and *FURIN* also have putative relationships with Ras family members (Wu et al., 2019; He et al., 2020) and were also only discovered by the FAB *t*-tests.

**Figure 4.**
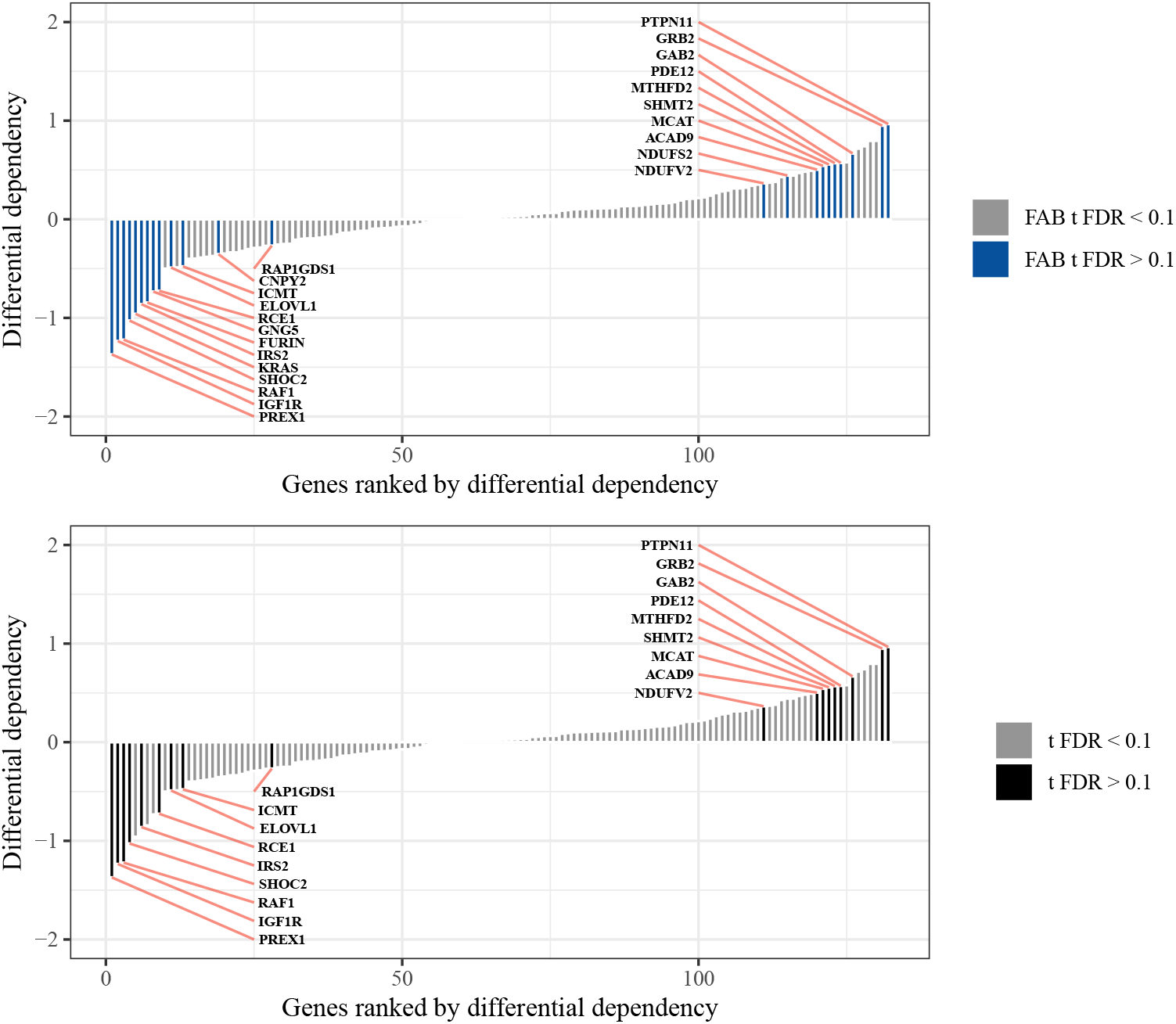
Results of differential dependency analysis between Ras-mutant and Ras-wild-type cancer cell lines from Wang et al. (2017). FAB *t*-tests (top) yield more discoveries than classical two-sided *t*-tests (bottom) at an FDR of 0.1.

Notably, this same testing scenario was considered in Bryan and Hoff (2022), where the authors used gene embeddings derived from a corpus of experimental data including CRISPR dependency scores, gene expression levels, copy number information, and mutation status of over 1,000 cancer cell lines. Comparing the results obtained here to the results from Bryan and Hoff (2022), we conclude that the embedding subspace determined by the GenePT embeddings was as useful for this task as the embedding subspace determined by the experimentally-derived gene embeddings.

### 2.5 Identifying genetic dependencies associated with metastasis

In another illustration of how gene embeddings derived from LLMs can guide statistical hypothesis testing in genomics studies, we consider multiple hypotheses concerning the presence or absence of an association between cancer metastasis and a binary categorization of dependency for each of several genes. In this type of association analysis, it is often the case that one would like to account for other factors, like the age or sex of a cell line donor, which might influence the association between metastatic status and genetic dependency. To test these hypotheses, we downloaded gene dependency scores for *n* = 861 cancer cells lines and *p* = 2906 genes along with the relevant age, sex, and metastatic status information from the Cancer Dependency Map. For this analysis, dependency on gene *j* was defined as having a dependency score above 0.5, and the number of genes we selected was based on the existence of at least 50 cancer cell lines with either a dependent / non-dependent status. The number of cell lines included was determined based on the availability of the age, sex, and metastasis information.

For each gene, we performed a logistic regression, treating dependency on the *j*th gene as the dependent variable, and age, sex, and metastatic status as the independent variables. The output of this analysis was a table of estimated regression coefficients and their approximate standard errors. The FAB *z*-test for logistic regression coefficients (Section 5.5) was then performed for each gene, as was the classical Wald *z*-test. In Figure 5 we display the results of this analysis. The left panel plots the observed regression coefficients for metastatic status versus their projection onto the 100-dimensional subspace spanned by the LLM-derived gene embeddings. The right panel displays the FAB and classical *p*-values after adjustment with the Benjamini-Hochberg procedure.

**Figure 5.**
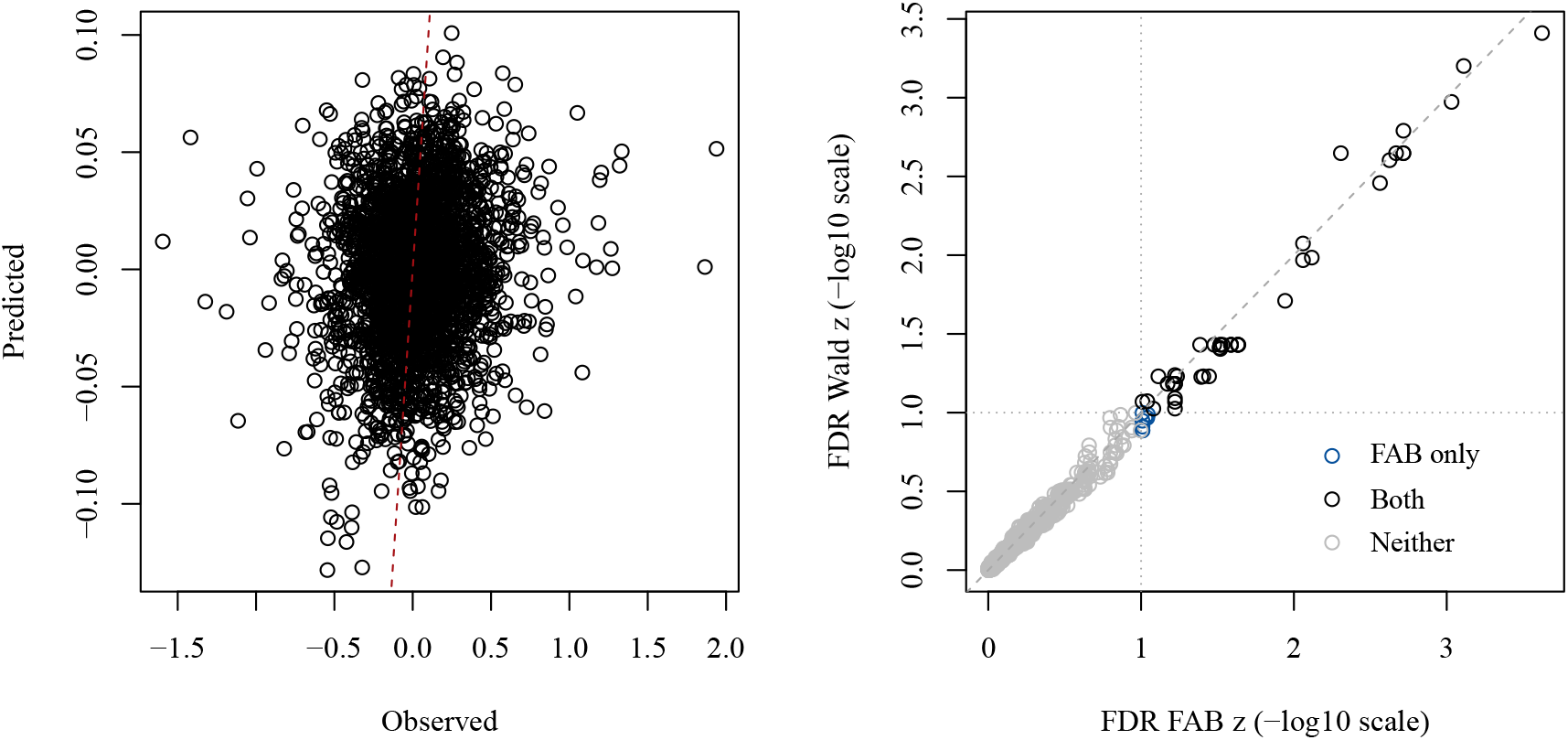
(left) Predicted regression coefficients based on projection to the 100-dimensional gene emedding subspace versus observed regression coefficients. (right) Adjusted *p*-values for the Wald *z*-tests versus those of the FAB *z*-tests. Discoveries made only by the FAB tests at an FDR of 0.1 are highlighted in blue.

Judging by the low correspondence between the projected and observed coefficients, the embedding subspace does not explain much of the variability among the regression coefficients. However, the FAB procedure adapts to this lack of fit. Referring back to Figure 2, the prior densities used for each FAB test tend to be centered near zero, and they tend to be diffuse, so the FAB test statistics are nearly the same as the classical Wald *z*-statistics. Importantly, though, they are not exactly the same. The slight correspondence between the projected and observed scores translates to several FAB tests with a modest, yet non-zero boost in power relative to their classical counterparts. As observed in the right panel of Figure 6, this means that at an FDR of 0.1, 45 discoveries were made only by the FAB *z*-tests, while no discoveries were missed, i.e. made only by the Wald *z*-tests.

**Figure 6.**
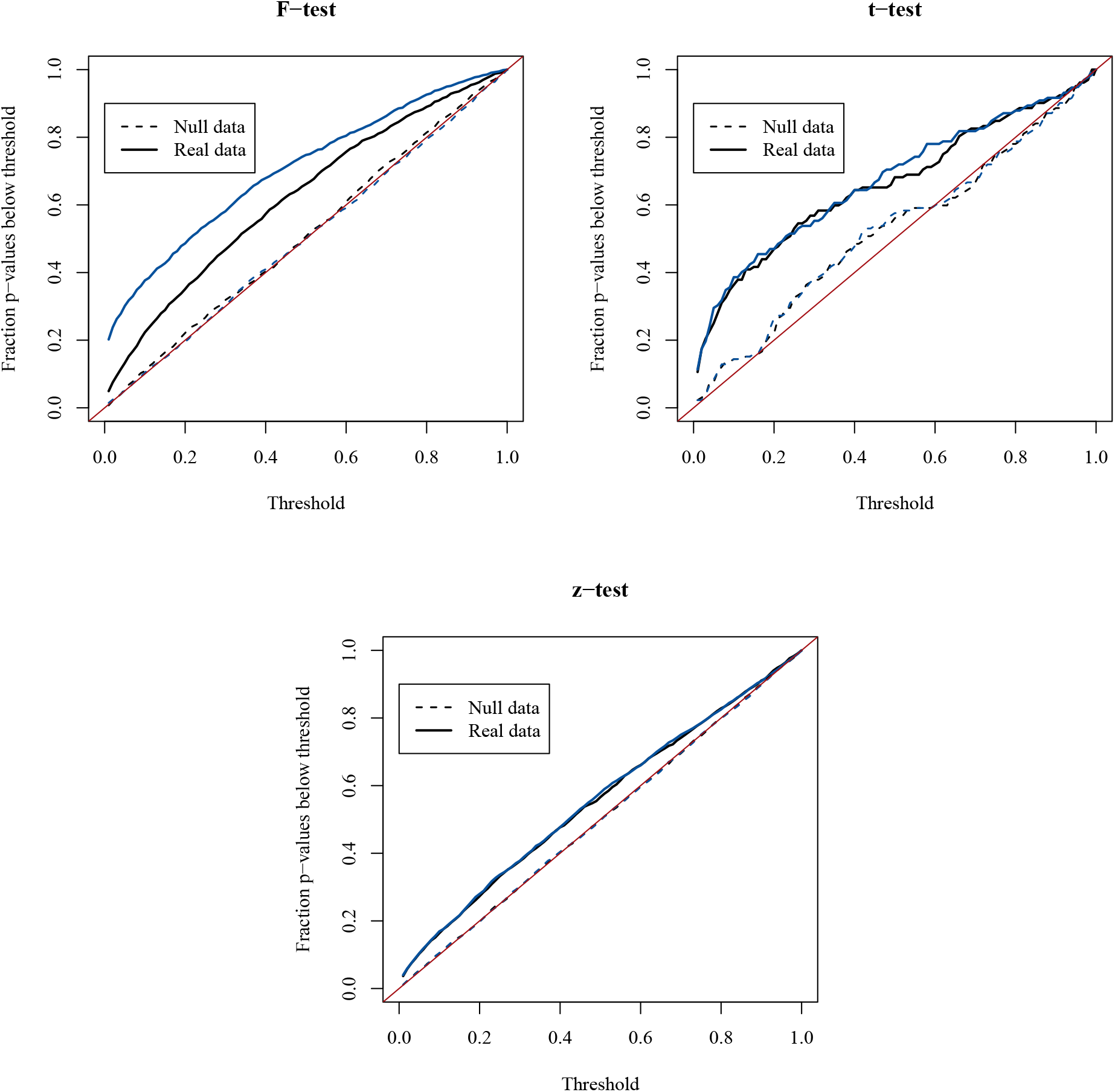
(left) the empirical CDFs of both the FAB and classical *p*-values show type I error control at the nominal level for data simulated from the null distribution, while the FAB *p*-values are stochastically smaller than the classical *p*-values for the drug viability data. (right) for the logistic regression analysis, the FAB *p*-values are stochastically smaller than the classical *p*-values, but less noticeably so. Still, the empirical CDFs of both the FAB and classical *p*-values show type I error control at the nominal level.

## 3 Simulation studies

### 3.1 Type I error of the FAB tests

The type I error guarantee of the three FAB tests we applied in Sections 2.2, 2.3, and 2.4 is that, under the null sampling model, the tests reject the null hypothesis with a pre-specified probability *α* due to the fact that the FAB *p*-values derived from these tests are theoretically guaranteed to be uniformly distributed between 0 and 1 under the null hypothesis. This is the same guarantee offered by classical statistical hypothesis tests. For the FAB test applied in Section 2.5, this guarantee holds only approximately, since the logistic regression coefficients are only approximately normally distributed. However, this normal approximation is also assumed by the classical Wald *z*-test. In summary, for all of the testing scenarios we consider in this article, the type I error guarantees of the FAB tests are the same as those of their classical counterparts.

The Methods section gives a theoretical explanation for why FAB tests behave like classical tests in terms of their type I error properties. Here, we provide empirical evidence that these properties hold for each of the four FAB tests we use in this article. In the simulations that follow, we generate data according to the probability distributions corresponding to the various null hypotheses described in the Methods section. For the FAB *F* -, *t*-, and *z*-tests, we set the problem dimensions *n, p*, and *d* equal to those found in the applications of Sections 2.3, 2.4, 2.5, respectively. For the FAB test of multivariate means, the problem dimensions are set to those described in the additional simulations of Section 3.2.

In Figure 6, we display the empirical CDFs of the FAB and classical *p*-values obtained from the simulated null data and those from the real data. Under the null distribution, both FAB and classical *p*-values should have a uniform distribution on the interval (0, 1), so their empirical CDFs should lie along the red line on the main diagonal of each plot. This is what we observe for all three tests, as evidenced by the dashed lines in each plot. Some deviation from the red line due to sampling error is expected since there are only finitely many hypotheses, and only one null simulation was run for each hypothesis. Particularly in the case of the AML analysis (*t*-test), there were only 132 hypotheses, so one would not expect perfect uniformity from the empirical CDF of these *p*-values. Indeed, the classical *p*-values display much the same deviation. In the metastatic association analysis (*z*-test), there were nearly 3000 hypotheses, so the slight deviation from uniformity observed in the null *p*-values may not be so easily attributed to sampling error. Rather, it is more likely due to the normal approximation used to motivate the *z*-test, which only holds asymptotically.

Once again, though, this is not unique to the FAB test. Since both the FAB and Wald tests use the normal approximation, comparable deviations from uniformity are observed for both sets of null *p*-values.

In Figure 7, we illustrate the behavior of the FAB test for high dimensional means on null data simulated from the scenarios described in Section 3.2. Each bar in the plot represents the proportion of rejections among 500 null simulations, where here the null sampling model refers to one for which there is no difference between the vector-valued means of two populations (in the notation of 3.2, ***δ*** = **0**). All versions of the FAB tests show type I error control at the nominal level *α* = 0.05, up to sampling error, as does the classical analog, which in this case is the random projection (RP) test. More details about the problem dimensions are given in the next section.

**Figure 7.**
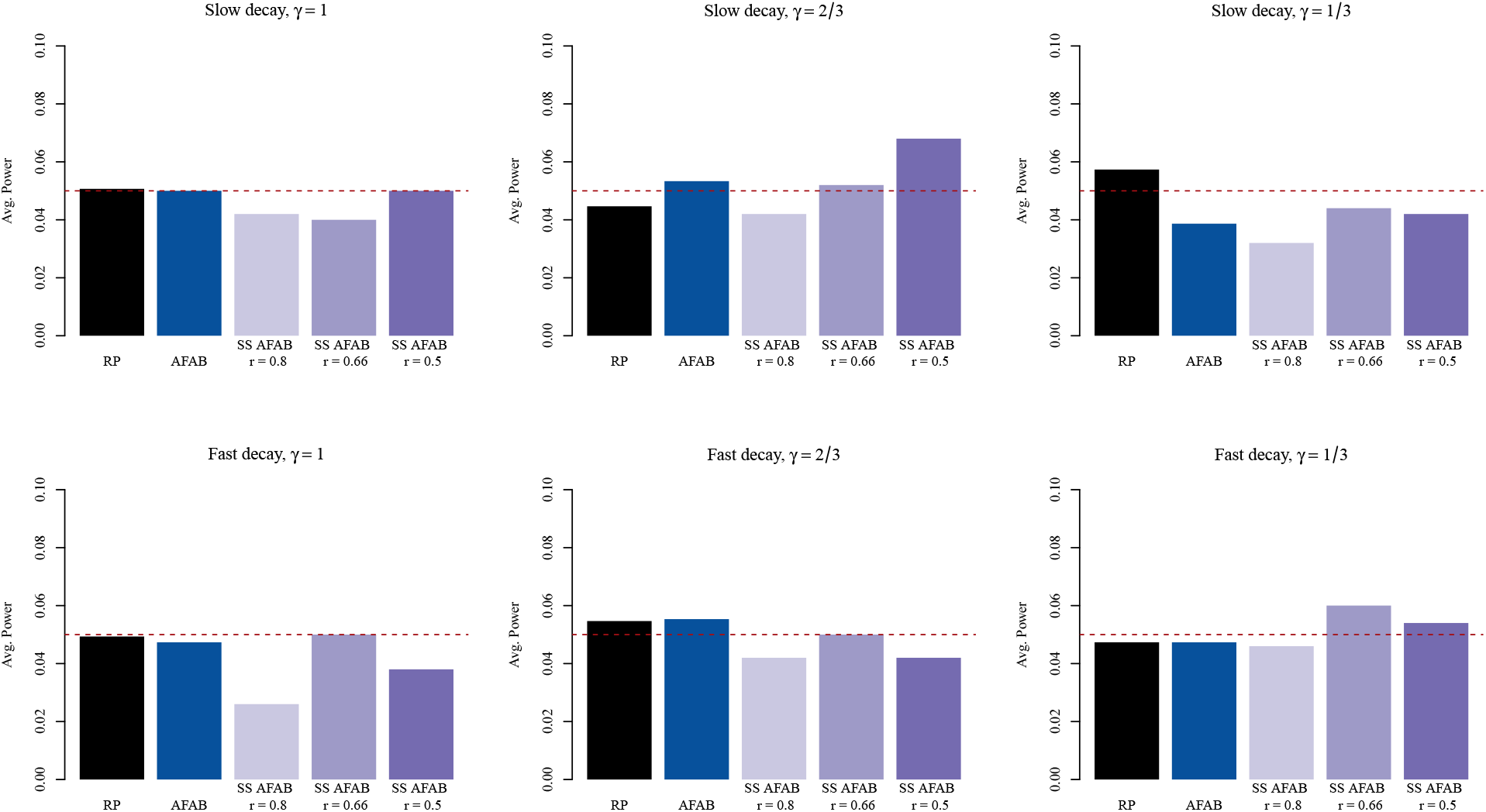
Average power for the FAB and RP tests under various simulation scenarios, all of which constitute null sampling models. Power under the null model is equivalent to type I error, and all tests exhibit type I error control at the nominal *α* = 0.05 level, up to sampling error.

### 3.2 Evaluation of the split-sample high dimensional means test

Here, we evaluate the AFAB test for detecting differences in multivariate means in a variety of simulation scenarios and compare its performance to that of the randomized projection (RP) test defined in Lopes et al. (2011) using *d* as the projection dimension. Letting *r* ∈ (0, 1), we also make a comparison between the naive AFAB test obtained by setting 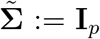 and a split-sample (SS) AFAB test, obtained by computing **y** and **S** using *rn*_1_ and *rn*_2_ observations, respectively, from each sample, and using a shrinkage estimate for 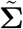 of the form

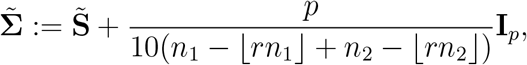

where 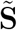 is the pooled covariance matrix obtained from the *n*_1_ − ⌊ *rn*_1_⌋ + *n*_2_ − ⌊ *rn*_2_ ⌋ observations not used to compute **y** and **S**. In our assessment of the SS AFAB test, we consider values of *r*∈ *{* 0.5, 0.66, 0.8*}* .

Prior to each simulation, we specify a matrix **E** ∈ ℝ^*p×d*^ with entries drawn independently from *N* (0, 1) and calculate the matrix of its left singular vectors **U**, which represent the gene embedding subspace used in the computation of the AFAB test statistics. We perform simulations for true error covariance matrices with two different rates of spectral decay by taking the eigenvalues of **Σ** to be equal to an equi-spaced sequence of *p* numbers between 0.1 and 1 raised to the power of 20 (fast decay) or 5 (slow decay). For both rates, the total variance is normalized so that trace(**Σ**) = 50. Within each of these scenarios, we evaluate the average power of each test across 15 random instances of **Σ** and ***δ*** = ***µ***_2_ − ***µ***_1_. The random covariance matrices are computed by setting the eigenvectors of **Σ** equal to a *p × p* orthogonal matrix simulated uniformly at random from 𝒱^*p*^. Given a noise variance *γ >* 0, random unit-norm signals are simulated relative to the embedding subspace according to the following generative process.

1. Simulate **z** ∼ *N*_*d*_(**0, I**_*d*_).
2. Simulate **v** ∼ *N*_*p*_(**Uz**, *γ***I**_*p*_)
3. Set ***δ*** = **v/**∥**v**∥.

For each of the 15 random instances of **Σ, *δ***, power is calculated as the proportion of rejected tests out of 100 realizations of the two samples 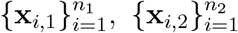 . In all simulations, *n*_1_ = *n*_2_ = 50, *p* = 200, and *d* = 10.

In Figure 8, we display the results of the simulation studies for the fast and slow decay scenarios and for each of three noise variance values *γ* ∈ *{* 1, 2*/*3, 1*/*3*}* . Overall, we observe that the AFAB tests have higher average power than the RP test. This is due to the fact that, even for the highest noise variance scenario (*γ* = 1) the simulated signals lie closer to the embedding subspace than would be expected if they were simulated uniformly at random from the unit sphere in ℝ ^*p*^. We also note that the average power of all tests considered is higher in the fast decay scenarios, which agrees with the results from Lopes et al. (2011).

**Figure 8.**
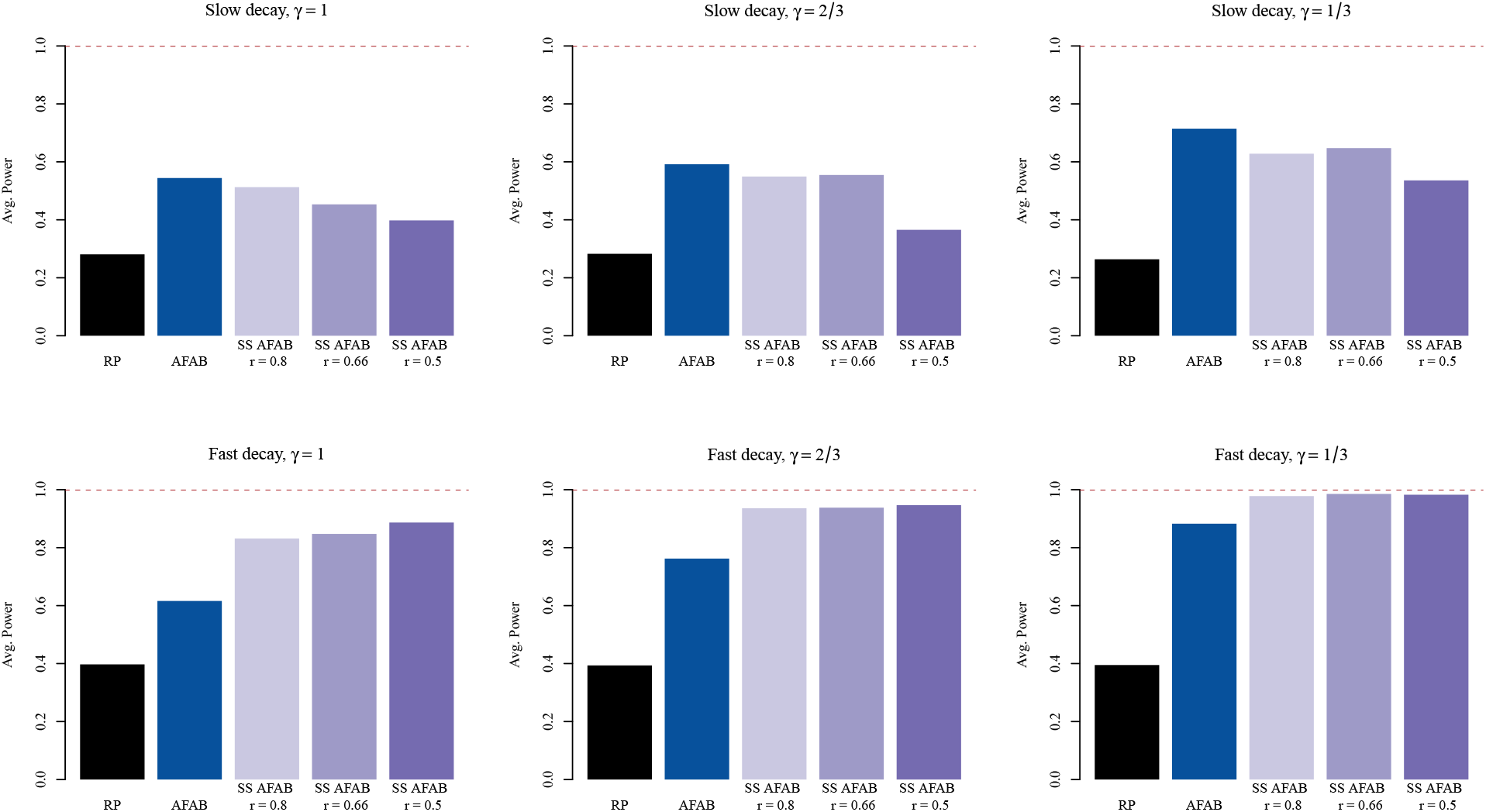
Average power for the FAB and RP tests under various simulation scenarios, all of which constitute sampling models with a unit-norm difference in high dimensional means. Spectral decay of the error covariance matrix increases from top to bottom, while average signal distance to the embedding subspace decreases from left to right.

In the slow decay scenarios, the naive AFAB test (denoted AFAB in Figure 8) outperforms the SS AFAB tests, regardless of the proportion of held-out samples used. This implies that whatever power was gained by using a data-driven value of 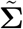 was offset by the loss in power due to having a lower sample size for the computation of **y** and **S**. However, in the fast decay scenarios, this tradeoff has the opposite result. Sample-splitting appears to confer a gain in average power, especially for the *γ* = 1 scenario. This suggests that the SS AFAB test may be most advantageous relative to AFAB in two-sample problems for which the high-dimensional covariance matrix is approximately low-rank and the true difference in means lies near to, but not in the embedding subspace.

## 4. Discussion

One of the challenges associated with performing statistical inference using information from LLMs is the “black-box” nature of that information. While the relationships between LLM-derived embeddings may be interpreted in terms of the entities they encode, the embeddings themselves have no straightforward interpretation in the absence of other data, which renders parametric models that use these embeddings as direct inputs uninterpretable. Our strategy for inference was to employ a FAB methodology for hypothesis testing, which uses the LLM information indirectly by focusing power for interpretable parametric hypotheses on regions of genomic space spanned by the LLM-derived gene embeddings. A key feature of the FAB hypothesis tests we proposed in this article is that they retain type I error control, regardless of the utility of the LLM-derived information for a particular set of biological hypotheses.

Interestingly, the real-data examples we examined suggest that that there is indeed useful information in the LLM-derived gene embeddings as pertains to a variety of genomics contexts. The increase in power we observed in the four applications of Section 2 can be attributed to the fact that biological signals arising in many genomics datasets lie close to the subspace spanned by the principal axes of the gene embeddings. The results presented in this article open several venues for further investigation.

While we employed gene embeddings from the GenePT model in this study, these embeddings are not the only possible representation of gene information. Investigating how embeddings derived from other models, such as GPT-4 or biology-specific models like BioBERT, impact hypothesis testing performance could provide additional insights into what types of LLM information are valuable for different hypothesis testing contexts. Additionally, assessing how embeddings trained on multimodal data (i.e. embeddings integrating both textual and experimental information) might further improve the power of FAB tests is a promising direction.

The FAB tests proposed in this article exploit linear relationships between the GenePT embeddings and biological signals. This is mostly due to the fact that FAB tests are derived within a classical statistical framework, and many of the multivariate statistical tests for which closed-form results are available involved linear models. Extending FAB methods to incorporate nonlinear functions of the gene embeddings as prior parameters could increase the flexibility of the FAB tests and potentially capture more complex relationships between genes that are encoded in the embeddings.

In part for computational reasons, this article discussed projection statistics resulting from limiting cases of FAB tests. However, in general the parameters *ν* and *γ* in Sections 5.2 and **??** can be chosen in a data-driven manner, as long as the data used to set the values of these parameters are independent of the data corresponding to the hypothesis being tested. Adaptive FAB tests like these may guard against a loss of power in biological contexts where the LLM gene embeddings are not informative.

## 5 Methods

To integrate LLM-derived information into the classical framework of statistical hypothesis testing, we use a methodology that is both frequentist and Bayesian (FAB). The four hypothesis tests we describe below are frequentist in the sense that they maintain a desired level *α*, while they are Bayesian in the sense that they are derived according to a criterion of maximal average power with respect to prior information. While such information may come from any source in general, here we show that the FAB approach provides a principled means to design test functions that are either optimal or approximately optimal with respect to the information derived from LLMs.

### 5.1 A statistical description of FAB hypothesis testing

We briefly review the general principle of FAB testing here before describing FAB tests for specific models. Suppose we observe an *m*-dimensional random vector **y** having a probability distribution *P*_***θ***_ with density *p*_***θ***_(**y**), both indexed by a *p*-dimensional parameter ***θ***. We wish to test the hypothesis *H* : ***θ*** = **0**, considering only those test functions *ϕ* : ℝ^*m*^→ {0, 1} that have type I error equal to a pre-specified level *α*. For a simple alternative ***θ*** = ***θ***_1_, the Neyman-Pearson lemma (NPL) provides the test function with maximum power:

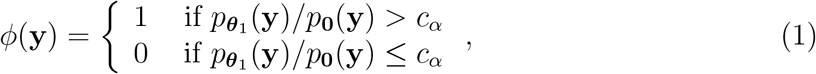

where *c*_*α*_ is a constant so that E[*ϕ*(**y**) | ***θ***] = *α* when ***θ*** = **0**. An appropriate test statistic for *H* may then be derived from the likelihood ratio 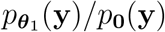.

The FAB approach derives test statistics for *H* according to a different kind of likelihood ratio, one which incorporates prior information about ***θ***. For our purposes, this prior information will be a function of LLM text embeddings **E**, so we represent this knowledge via a probability distribution having density *π*_**E**_. On average with respect to *π*_**E**_, a test function *ϕ* has power

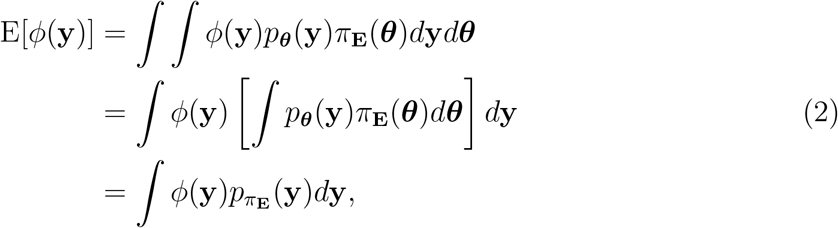

where 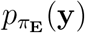 is the marginal density of **y** under *π*_**E**_. Invoking NPL again, the level-*α* test that maximizes (2) rejects *H* when the likelihood ratio 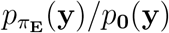 is greater than its 1 − *α* quantile assuming the null distribution **y** ∼ *P*_**0**_. Such a test is FAB because among all frequentist level-*α* tests for *H* it is the Bayes-optimal test with respect to the LLM-derived information in *π*_**E**_. As it is often more convenient to work in terms of the log-likelihood ratio 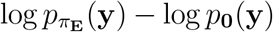 instead of the likelihood ratio itself, we do so for the tests derived in the following sections.

### 5.2 A FAB test for multivariate means

In this section we develop a FAB test for the two-sample multivariate mean hypothesis testing problem that uses prior information derived from LLM text embeddings. Our primary interest lies in applications to perturbation screens in genomics, in which a high-dimensional phenotype such as mRNA expression for thousands of genes is recorded for both treated and untreated samples of cells. In the context of a high-throughput screen where treatments consist of small molecules or gene edits of unknown effect, one question of interest is whether the average gene expression profiles between the treated and untreated cell populations differ at all. Answering this question distinguishes perturbations that show evidence of bioactivity from those that appear to be inert.

To motivate the FAB test for a single perturbation, we assume the sampling model

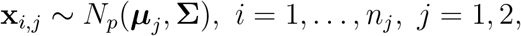

where **x**_*i,j*_ represents the *i*th gene expression profile from treatment category *j*, and *j* = 1 corresponds to “untreated.” The hypothesis we want to test is *H* : ***µ***_1_ = ***µ***_2_. For now, let **Σ** be known, so that 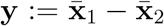 is sufficient for ***θ*** := ***µ***_1_ − ***µ***_2_, and we may simplify the problem to testing *H* : ***θ*** = **0** with

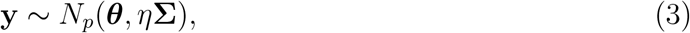

and *η* = 1*/n*_1_ + 1*/n*_2_. We encode the belief that biologically meaningful differences in average gene expression profiles between treated and untreated cell populations will be close to the principal subspace described by the LLM embeddings using the prior distribution

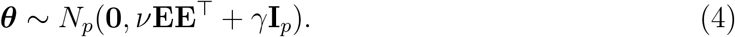

Now, we can directly apply the reasoning from Section 5.1 (with *m* = *p*) to obtain an oracle FAB test for *H*.

#### Proposition 1.

*The level-α FAB test of H corresponding to sampling model* (3) *and prior distribution* (4) *rejects when the test statistic*

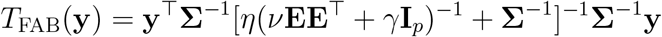

*exceeds its* 1 − *α quantile under H*.

The null distribution of *T*_FAB_(**y**) is that of a weighted sum of *p* independent 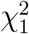 random variables, where the weights are equal to 1*/*(*ηλ*_*j*_ + 1) for *j* = 1, …, *p*, and *λ*_*j*_ is equal to the *j*th eigenvalue of **Σ**^1*/*2^(*ν***EE**^⊤^ + *γ***I**_*p*_)^−1^**Σ**^1*/*2^. If **Σ** is known, then the quantiles of this distribution may be approximated with high precision using Monte Carlo.

In practical settings, **Σ** will not be known. To arrive at a computable approximation to *T*_FAB_, we first consider a limiting case of (4), which occurs when *γ* → 0 and *ν* → ∞, leading to a diffuse prior distribution with all of its mass on the embedding subspace. The FAB test statistic corresponding to this limiting prior distribution, termed *T*_LFAB_(**y**), is given in the following proposition.

#### Proposition 2.

*Let* **A** = **Σ**^−1^**E**. *Then*

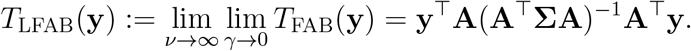

The expression for *T*_LFAB_(**y**) is written to emphasize the fact that it is a projection statistic in the sense that it is invariant to right multiplications of the form **A** *1*→ **AG** for invertible *d × d* matrices **G**. Therefore, *T*_LFAB_(**y**) depends only on the column space of **A**, which is determined in part by the column space of the LLM embeddings. Tests based on projection statistics have been shown to have favorable power relative to other tests for differences in high dimensional means (Lopes et al., 2011; Jacob et al., 2012), which motivates this choice of prior distribution.

Approximating *T*_LFAB_(**y**) yields an embedding-projected version of Hotelling’s *T* ^2^ statistic, which is only roughly Bayes-optimal with respect to the limiting case of (4), but which has a tractable finite-sample null distribution. Let

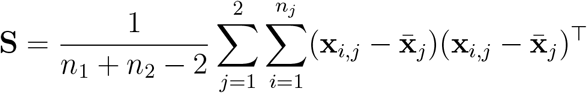

be the pooled sample covariance matrix of the gene expression profiles, and suppose further that an independent estimate 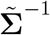 of **Σ**^−1^ is available. Then our proposed approximate FAB (AFAB) statistic is

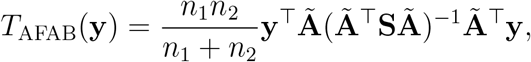

Where 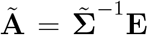. By the mutual independence of **y, Ã**, and **S**, the null distribution of *T*_AFAB_(**y**) can be characterized by

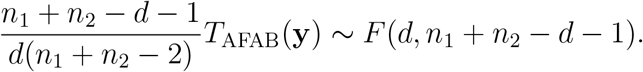

Therefore, a level-*α* test is to reject *H* when *T*_AFAB_(**y**) exceeds 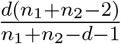 times the 1 − *α* quantile of the *F* (*d, n*_1_+ *n*_2_ − *d* − 1) distribution.

#### 5.2.1 Choice of 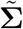

As long as 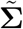 is independent of **y** and **S**, the test based on *T*_AFAB_ will have level-*α* regardless of the particular choice of 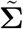 . The power of the AFAB test, though, will vary according to the column space of 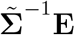 **E**. If the naive choice of 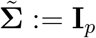 is made, *T*_AFAB_ is equivalent to a non-randomized version of the Lopes et al. (2011) statistic, where the projection matrix is **E** rather than a matrix of entries drawn independently from *N* (0, 1). Alternatively, one can take a split-sample approach, reserving a fraction of the **x**_*i,j*_’s to estimate 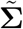 independently of **S**. The following proposition summarizes why it may be worthwhile to find a non-naïve 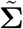 that approximates the true **Σ**.

##### Proposition 3.

*Let* 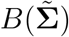 *denote the power of the AFAB test as a function of* 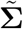 . *Then for any* ***δ*** = ***µ***_2_ − ***µ***_1_ *in the column space of* **E**, *it holds that*

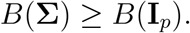

This result conforms with our previous reasoning about the maximal power property of exact FAB tests, and it suggests that using a 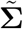 that closely approximates the true **Σ** could result in a more powerful test than naively setting 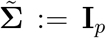. However, the split-sample approach will reduce the number of observations used to compute **S**, thereby reducing the power of the corresponding test. We explore the trade-off induced by the split-sample approach in the simulation of Section 3.2.

### 5.3 A FAB *F* -test for linear hypotheses

In other high-throughput screening contexts, a perturbation is applied to a panel of diverse cancer cell lines, and a univariate phenotype such as post-perturbation cell viability is recorded for each cell line. For example, the data collected by Corsello et al. (2020) consist of post-treatment cell viability scores for 905 cancer cell lines for each of 1518 drugs with both oncology and non-oncology indications. In this circumstance, it is of interest to determine which drugs produce a biologically meaningful cancer-killing pattern, especially as the killing pattern relates to genomic variables such as baseline mRNA expression.

Here we describe how LLM gene embeddings can be incorporated into tests of linear hypotheses that relate cell viability to baseline mRNA expression. To do so we adopt the FAB method of McCormack and Hoff (2023), which modifies the classical *F* -test to incorporate prior information. A direct application of this method would share information across perturbations to specify the FAB prior, potentially biasing the FAB test results towards mechanisms of action that are over-represented in the perturbation library. Instead, we treat perturbations independently, and we use the LLM gene embeddings to boost power near biologically salient subspaces of ℝ^*p*^.

For a single perturbation, we assume that a vector of *n* viability scores follows the sampling model

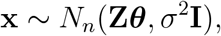

where **Z** ∈ ℝ^*n×p*^ is a matrix of mRNA expression values for *n* cell lines and *p* genes. The hypothesis *H* : ***θ*** = **0** corresponds to a perturbation for which the pattern of viability scores shows no discernible relationship with any linear combination of gene expression profiles. Following McCormack and Hoff (2023), we consider scale-invariant tests based on the self-normalized viability profile **y** = **x***/*∥ **x**∥, which has an angular Gaussian distribution on the *n* − 1 unit sphere 𝒮^*n*−1^. In contrast to the previous section, this allows us to remove dependence on the unknown *σ*^2^ from the log likelihood ratio defining the FAB test statistic because the distribution of **y** under *H* is the uniform distribution on 𝒮^*n*−1^. Specifying a normal prior distribution on ***θ*** that places mass near the embedding principal subspace

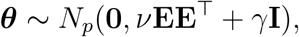

and placing a point-mass prior distribution on 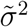 for *σ*^2^, the resulting FAB statistic is

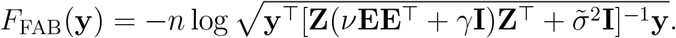

While the null distribution of *F*_FAB_(**y**) does not have a simple form, its quantiles may be approximated with Monte Carlo by simulating **x**^(1)^, …, **x**^(*S*)^ ∼ i.i.d. *N*_*n*_(**0, I**), and then evaluating the quantiles of 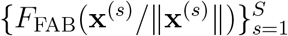 . A FAB test with level arbitrarily close to *α* may then be performed by rejecting *H* when *F*_FAB_(**y**) exceeds the 1 − *α* quantile of this empirical null distribution for *S* large.

For embedding dimension *d < n*, we can further simplify the FAB statistic by considering the limiting case *ν*→ ∞, *γ* → 0 as we did in the previous section. Doing so, we obtain another embedding-projected statistic, which approximates the FAB *F* statistic when *ν* is very large relative to *γ* and 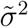.

#### Proposition 4.

*For d < n, the limiting FAB F -test as ν* → ∞, *γ* → 0 *rejects H* : ***θ*** = **0** *when the classical F -test rejects H* : ***β*** = **0** *under the sampling model*

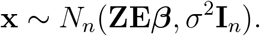

In Section 2.3, we apply this limiting-case FAB test to a collection of linear hypotheses concerning the Corsello et al. (2020) viability scores.

### 5.4 FAB *t*-tests for multiple scalar hypotheses

In the previous two sections, we discussed FAB tests for *p*-dimensional hypotheses, using information in LLM embeddings to increase power in certain directions of ℝ ^*p*^ where we expect biologically meaningful signals to reside. Here we describe how the FAB principle may be used to test several 1-dimensional hypotheses as may arise when performing a differential expression analysis between two samples of biological specimens. In this case, the embeddings provide a means to borrow information across the independent experiments providing the data for each hypothesis, rather than providing a subspace on which to concentrate power for a multivariate hypothesis.

Assume that a transcriptomic measurement is recorded for each of *p* genes in each of 2 treatment categories with *n*_1_, *n*_2_ replicate experiments per category. A sampling model for the measurement corresponding to the *i*th replicate of the *j*th gene in the *k*th treatment category is

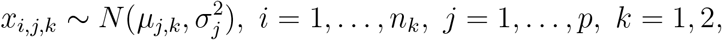

where it is assumed that, conditional on the population-level means *µ*_*j,k*_, the *x*_*i,j,k*_ are mutually independent. Difference scores are obtained by taking the difference between the average measurement in the two treatment categories, leading to the condensed sampling model

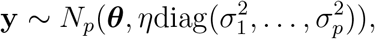

where 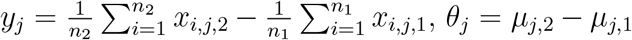, and *η* = 1*/n*_1_ + 1*/n*_2_. In order to identify which genes exhibit differential transcriptomic activity, it is of interest to test the hypothesis *H*_*j*_ : *θ*_*j*_ = 0 for each *j* ∈ {1, …, *p*} .

In this scenario, we consider FAB *t*-tests informed by LLM-derived gene embeddings based on normal prior distributions

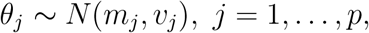

where *m*_*j*_ and *v*_*j*_ are chosen to reflect the prior belief that the vector of true differences ***θ*** will be close to the principal embedding subspace. As long as *m*_*j*_ and *v*_*j*_ are independent of *y*_*j*_, an approximately Bayes-optimal test for *H*_*j*_ may be based on the *p*-value

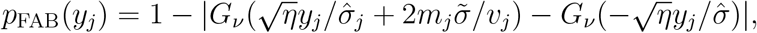

where *G*_*ν*_ is the CDF of the *t*-distribution with *ν* degrees of freedom, 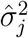 is the pooled sample variance of the transcriptomic measurements for gene *j*, and 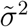 is an independent estimate of *σ*^2^ (Hoff, 2020). LLM-informed values of *m*_*j*_ and *v*_*j*_ may then be obtained by computing the posterior expectation and variance, respectively, of *θ*_*j*_ with respect to the Bayesian model

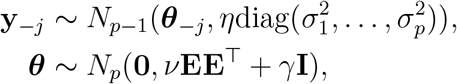

where **y** _−*j*_ denotes the vector consisting of all but the *j*th difference score. This method for FAB *t*-testing is analogous to that in Bryan and Hoff (2022), where more details, including those regarding adaptive selection of the prior parameters *ν* and *γ* and computation of 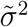, may be found.

### 5.5. FAB *z*-tests for logistic regression coefficients

In previous subsections, we have described tests that are approximately Bayes-optimal in settings where the sampling model holds exactly. Here, we describe how the FAB principle may be used to conduct tests that are exactly Bayes-optimal in a setting where the assumed sampling model holds approximately. Specifically, we consider testing multiple scalar hypotheses, where each test statistic is related to a Wald *z*-score derived from the estimated coefficient in a logistic regression.

Suppose that *x*_*i,j*_ is a binary random variable, perhaps representing dependency status for cell type *i* and gene *j*, which is equal to 1 when the dependency is present and 0 when it is not. A common sampling mechanism for *x*_*i,j*_ is specified by the logistic regression model

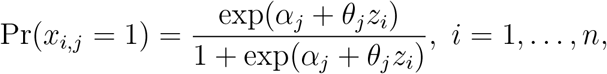

where *z*_*i*_ is the value of a covariate of interest for cell type *i, α*_*j*_ is the baseline log-odds of dependency for gene *j*, and *β*_*j*_ is a regression coefficient representing the change in log-odds of dependency of gene *j* per one-unit change in the value of the covariate. To assess whether the covariate has an association with the dependency status of genes *j* = 1, …, *p*, one may evaluate *H*_*j*_ : *θ*_*j*_ = 0 for each gene.

While the distribution of the maximum likelihood estimate (MLE) of *θ*_*j*_ cannot be written in closed form, results from statistical theory say that it is approximately normal for *n* sufficiently large, with a mean and variance that both depend on *θ*_*j*_. By related arguments, the Wald statistic

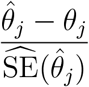

has an approximate standard normal distribution for large *n*, where 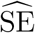 denotes a plug-in approximation to the asymptotic standard error of the MLE 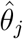, which can be obtained from the inverse information matrix of *θ*_*j*_ in the logistic regression model. This motivates the following approximate sampling model for the vector of estimated regression coefficients obtained via maximum likelihood

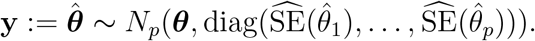

FAB tests that incorporate LLM-derived information can be based off of the FAB *p*-values described in the previous section, which use the prior distribution ***θ*** ∼ *N*_*p*_(**0**, *ν***EE**^⊤^ + *γ***I**). The only modification needed is that the CDF of the *t*-distribution with *ν* degrees of freedom is replaced with the standard normal CDF, this being due to the fact that under the approximate sampling model the standard errors are taken to be fixed and known rather than quantities that need to be estimated.

## A Data and Embeddings

The datasets used for the genomic experiments are the Broad Dependency Map dataset (DepMap, 2024; DepMap and Kocak, 2024), and the Perturb-seq dataset (Replogle et al., 2022) deposited at https://plus.figshare.com/articles/dataset/_Mapping_information-rich_genotype-phenotype_landscapes_with_genome-scale_Perturb-seq_Replogle_et_al_2022_processed_Perturb-seq_datasets/20029387. The LLM embeddings used in all experiments are from GenePT (Chen and Zou, 2023), and were derived from the text-based NCBI (Schoch et al., 2020) gene descriptions using GPT-3.5 publically available through https://github.com/yiqunchen/GenePT and deposited at https://doi.org/10.5281/zenodo.10833191.

## B Proofs

### B.1 Proof of Proposition 1

*Proof*. Given sampling model

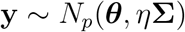

and prior distribution

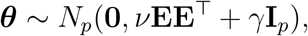

the level-*α* FAB test for *H* : ***θ*** = **0** rejects when the log-likelihood ratio

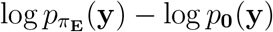

exceeds its 1 − *α* quantile under *H*. Removing terms that do not depend on **y**, the log-likelihood ratio is proportional to

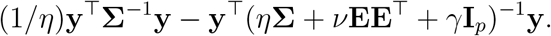

Now apply the Woodbury matrix inversion identity (Woodbury, 1950) to cancel the term (1*/η*)**y**^⊤^**Σ**^−1^**y** and obtain

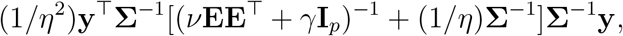

which is proportional to

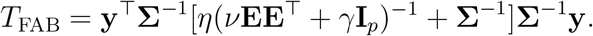

### B.2 Proof of Proposition 2

*Proof*. From above, the relevant portion of the log-likelihood ratio is

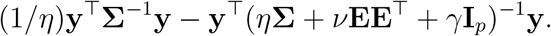

Set **Ψ** := *η***Σ** + *γ***I**_*p*_, and apply the Woodbury matrix identity to a different grouping of terms than above to obtain

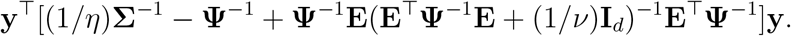

Now, lim_*γ*→0_ **Ψ**^−1^ = (1*/η*)**Σ**^−1^, so the limit as *γ* → 0 of the expression above is

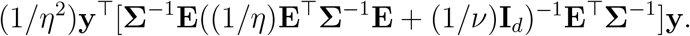

Taking the limit of this expression as *ν* → ∞ yields

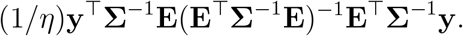

Making the substitution **A** = **Σ**^−1^**E** and comparing the above to *T*_FAB_ completes the proof.

### B.3 Proof of Proposition 3

*Proof*. As a function of 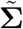, the power of the AFAB test is determined by the non-central *F* (*d, n*_1_ + *n*_2_ − *d* − 1) distribution, and increases monotonically with the non-centrality parameter

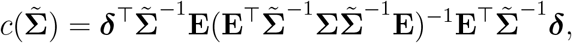

where ***δ*** = ***µ***_2_ − ***µ***_1_. To show that the power of an oracle version of the AFAB test has greater power than the AFAB test based on 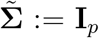 for any ***δ*** in the column space of **E**, we will show that *c*(**Σ**) ≥ *c*(**I**_**p**_) for any such ***δ***.

An oracle version of the AFAB test would have non-centrality parameter

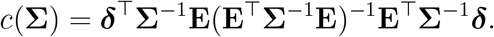

Because of the invariance of the AFAB statistic, without loss of generality we can assume **E** is orthogonal. For any ***δ*** in the column space of **E**, we may write ***δ*** = **Ed** for some **d** ∈ ℝ^*d*^. Thus, the non-centrality parameter of the oracle AFAB test is

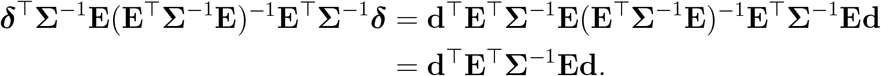

Compare this to the power of the AFAB test setting 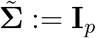, which is

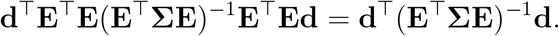

By the Gauss-Markov Theorem (David, 1938), we have

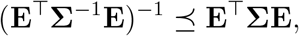

where ⪯ denotes the Loewner partial order. Therefore,

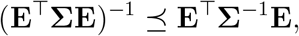

which shows that *c*(**Σ**) ≥ *c*(**I**_*p*_).

### B.4 Proof of Proposition 4

*Proof*. Let *d < n*. The FAB *F* -test is based on the test statistic

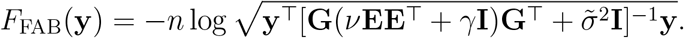

When *γ* → 0, we have

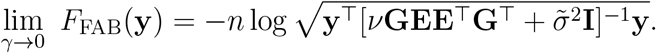

Let **Q, Λ** be, respectively, the left singular vectors and singular values of the matrix **GE**. Then

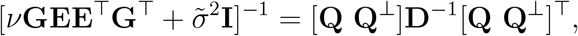

where **Q**^⊥^ is an orthogonal basis for the column space of **I**_*n*_ − **QQ**^⊤^, and **D** is a diagonal matrix with entries 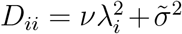 for *i* = 1, …, *d* and 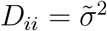 for *i* = *d* + 1, …, *n*. Taking *ν* → ∞, we have

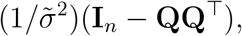

so

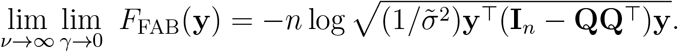

Now, both this limiting statistic and the *F* -statistic based on the sampling model

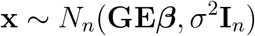

are strictly increasing functions of the quantity **y**^⊤^**QQ**^⊤^**y**. Therefore, a level-*α* test based on lim_*ν*→∞_ lim_*γ*_→_0_ *F*_FAB_(**y**) rejects when the level-*α F* -test for *H* : ***β*** = **0** under the sampling model above rejects.

